# Generation and utilization of a HEK-293T murine GM-CSF expressing cell line

**DOI:** 10.1101/2020.06.18.160366

**Authors:** Elektra K. Robinson, Sergio Covarrubias, Simon Zhou, Susan Carpenter

## Abstract

Macrophages and dendritic cells (DCs) are innate immune cells that play a key role in defense against pathogens. *In vitro* cultures of bone marrow-derived macrophages (BMDMs) and dendritic cells (BMDCs) are well-established and valuable methods for immunological studies. Typically, commercially available recombinant GMCSF is utilized to generate BMDCs and is also used to culture alveolar macrophages. We have generated a new HEK-293T cell line expressing murine GM-CSF that secretes high levels of GM-CSF (∼180ng/ml) into complete media as an alternative to commercial GM-CSF. Differentiation of dendritic cells and expression of various markers were kinetically assessed using the GM-CSF HEK293T cell line, termed supGM-CSF and compared directly to purified commercial GMCSF. After 7-9 days of cell culture the supGM-CSF yielded twice as many viable cells compared to the commercial purified GM-CSF. In addition to differentiating BMDCs, the supGM-CSF can be utilized to culture alveolar macrophages without an altering inflammatory activation cascade. Collectively, our results show that supernatant from our GM-CSF HEK293T cell line supports the differentiation of mouse BMDCs or alveolar macrophage culturing, providing an economical alternative to purified GM-CSF.

## Introduction

Colony-stimulating factors (CSF) including macrophage colony-stimulating factor (M-CSF), granulocyte colony-stimulating factor (G-CSF), and granulocyte-macrophage colony stimulating factor (GM-CSF) are crucial for survival, proliferation, differentiation and functional activation of hematopoietic cells, including macrophages and dendritic cells (DCs) (*1*). Macrophages and DCs are innate immune cells found in tissues and lymphoid organs that play a key role in defense against pathogens (*2*). While there are a multitude of macrophage and dendritic cell subsets, GM-CSF is critical for the development of conventional dendritic cells (cDCs) and alveolar macrophages (AMs) (*2*). Due to cell number limitations from harvesting cDCs and AMs directly from mice, well-established *in vitro* culturing of bone marrow and bronchoalveolar lavage fluid for dendritic cells and alveolar macrophages, respectively, using GM-CSF have become invaluable for immunological and molecular biology studies (*2*). This has led to the use of CSF proteins in the purified form, as well as to the generation of recombinant cell lines that secrete the desired protein in the supernatant for cost efficiency (*3–6*). One of the most widely adapted cell lines, utilized for differentiating and culturing murine bone-marrow derived macrophages (BMDMs) is the NCTC clone 929 strain L line, also known as L929 (*7*). Supernatant from cultured L929 can be utilized, in lieu of purified M-CSF, to culture and differentiated BMDMs because it secretes murine M-CSF (*7*).

The Human embryonic kidney 293T (HEK293T) cell line is the ideal choice for expressing a CSF protein for differentiating primary immune cells from mouse bone marrow as it has been shown not to express innate immune pattern recognition receptors or naturally secrete immune-related cytokines (*8, 9*). This ensures that it is predominantly only your protein of choice being expressed and that there is no inadvertent activation of the inflammatory cascade. Previously, murine GM-CSF has been cloned into J558L, a mouse B myeloma cell line (*4*), which is known to express cytokines, including IL-10 (*10*). Using J558L can therefore alter the results of *in vitro* experiments, by activating anti-inflammatory pathways when cultured with BMDCs or alveolar macrophages.

In this study we constructed HEK293T cell lines stably expressing and secreting murine GM-CSF. We utilized GM-CSF to generate and culture BMDCs and alveolar macrophages (AMs). HEK293T cells have no expression of human GM-CSF and the constructed cell lines thus express only the stably transfected gene of murine GM-CSF. We have found that our line is very stable, producing GM-CSF at a concentration of ∼200ng/ml even following freeze thaw cycles of the line. We differentiated BMDCs and cultured AMs using our supGM-CSF and compared them to commercially available purified GM-CSF (pGM-CSF) and found that our supGM-CSF yields a higher number of cells, purity of DCs is not altered, and have an intact immune signaling cascades.

## Materials and Methods

### Cloning Strategy of mGM-CSF

The mGM-CSF gene, including PspXI and NotI restriction sites, was amplified from the pCR3.1-mGM-CSF vector (Addgene, 74465). The PCR was set up with:

25 μl 2X Phusion High-Fidelity PCR Master Mix (Thermo Scientific), 1 μl PspXI_hMCSF_fwd (20mM): TCCGCTCGAGCCACCATGTGGCTGCAGAATTTACTTTTCC, 1 μl NotI_hMCSF_rev (20mM): GACGCGGCCGCTCATTTTTGGCCTGGTTTTTTGC, 1 μl pCR3.1-mGM-CSF (20ng) and 22 μl DEP-C nuclease-free water. PCR program: 95°C 3 minutes, 35 cycles of 95°C 30 sec, 60°C 30 sec, and 72°C 1 min, and end the PCR with 72°C and 12°C hold. The PCR product was purified using the PCR Clean-up Kit (Macherey-Nagel) and was subsequently digested with PspXI (New England BioLabs) and NotI-HF (New England BioLabs) using recommended digestion conditions (https://nebcloner.neb.com/#!/redigest) and was cloned into our custom 681 bidirectional lentiviral vector (sequence in supplemental). Sequence was confirmed by Sequetech (Sanger) sequencing.

### Lentivirus Generation

HEK-293T cells (4e5 cells/well) were plated onto a 6-well plate (353046, Corning) with complete DMEM [10% heat-inactivated FCS (Gibco, 26140-079), 100μg/ml penicillin (Thermo, 15140122), and 100μg/ml streptomycin (Thermo, 15140122)]. 24hr later, 500 ng of 681-mGM-CSF or empty vector control, 250ng psPAX2 (Addgene, 12260), and 250ng pMD2.G (Addgene, 12259) were mixed in 200μl of serum-free Opti-MEM (Gibco) and 5μl Lipofectamine 2000 (Thermo Fisher) was added to mix and incubated for 20 minutes at room temperature. Transfection reaction was added to HEK-293T cells and allowed to transfect for 72 hours, and supernatant was harvested, passed through 0.45μm filters (Millipore, Stericup), and aliquots were stored at -80°C.

### Construction of GM-CSF-producing HEK-293T Cells

HEK-293T cells were transduced 2 days with 200ul of lentivirus per 1e5 cells. 48hrs after infection, the HEK-293T cells were selected with puromycin (2ug/ml) for >3 days, monitoring viability and increase of mCherry expression using FACS (Attune NxT Flow Cytometer). Once mCherry expression exceeded 85%, puromycin was removed, and cell-line was expanded and used to produce GM-CSF. A detailed method for GM-CSF supernatant production is described in extended methods.

### Enrichment of DCs and Macrophages from BM and in-vitro stimulation

Bone marrow (BM) cells were harvested from the femurs and tibia of wild-type C57BL/6 between 6- and 18-weeks old mice and depleted of erythrocytes. 1e6-3e6 BM cells were plated per well of a 6-well tissue culture plate (353046, Corning) in complete DMEM supplemented with either 10% M-CSF from L929 cells, 5-10% supernatant, or 10-25ng/ml of recombinant mGM-CSF [PMC2015, Thermo Fisher]. Media was replaced on day 3 and every 2 days henceforward. Cells were scraped and moved onto a larger plate as they proliferated. After 7 to 14 days of enrichment, cells were stimulated by adding 200ug/ml LPS (Sigma, L2630-10MG) to the media and harvested for analysis after 0, 6, or 24 hours of stimulation. Total cell count was determined by staining cells with trypan blue and using a Light Microscope and hemocytometer.

### Surface Staining for Flow Cytometry

Stimulated BMDCs and BMDMs were harvested and centrifuged at 300 g for 5 minutes. The pellets were resuspended in 100μl sorting media (2% FCS, 5 mM EDTA, 1X PBS). Each sample was blocked with 100μl of Fc block CD16/CD32 (BD Biosciences) diluted 1:250 in sorting media and incubated at room temperature for 15 minutes. Antibodies and viability dye were then used to stain the samples for 30 minutes on ice in the dark. Antibodies and dye: APC-eFluor 780 anti-CD45 (Invitrogen, 47-0451-82), Alexa-Fluor 700 anti-CD11c (BD Pharmingen, 560583), FITC anti-CD11b (Thermo Fisher, MA1-10081), PE-eFluor 610 anti-F4/80 (eBioScience, 61-4801-80), PE anti-MHC Class II (BD Pharmingen, 562010), and Fixable Aqua Dead Cell Stain (Thermo Fisher, L34957). Cells were washed twice in 1 ml sorting media, spinning at 300 g for 5 minutes. The resulting pellet was resuspended in 200μl of sorting media and analyzed by FACS (Attune NxT Flow Cytometer).

### Inflammasome Activation Assay and ELISA

supGM-CSF (25ng/ml) or recGM-CSF (25ng/ml) differentiated BMDCs were plated at 2e5cells/well in a 96 well plate with complete DMEM media. Cells were first primed with 200ng/ml LPS for 3h prior to treatment with different agents. Poly(dA:dT) DNA (dsDNA mimic) was transfected using Lipofectamine 2000 at a concentration of 1.5μg/ml, 6hrs prior to harvesting supernatant. The supernatant was harvested using a p300 multichannel pipette and stored at -80°C. Commercial ELISAs were used to measure the following analytes in triplicate mice following the manufacturer’s instructions: IL-6 (mouse IL-6 DuoSet, R&D Systems DY406) and IL-1β (Mouse IL-1β/IL-1F2 R&D Systems DY401-05).

### BALF Harvest and Alveolar Macrophage Culturing

Bronchoalveolar Lavage Fluid (BALF) was harvested as previously stated by Cloonan *et al*. (PMID:26752519). 40 mice were euthanized by CO2 narcosis, the tracheas cannulated, and the lungs lavaged with 0.5-ml increments of ice-cold PBS eight times (4 ml total), samples were combined in 50ml conical tubes. BALF was centrifuged at 500*g* for 5 min. 1 ml red blood cell lysis buffer (Sigma-Aldrich) was added to the cell pellet and left on ice for 5 min followed by centrifugation at 500*g* for 5 min. The cell pellet was resuspended in 500μl PBS, and leukocytes were counted using a hemocytometer. Specifically, 10μl was removed for cell counting (performed in triplicate) using a hemocytometer. Cells were plated in sterile 12 well plates at 5e5/well (total of 8 wells) and use complete DMEM with 25ng/ml supGM-CSF.

### Culturing of Alveolar Macrophages

24hrs post-BALF isolation, media was removed and fresh complete DMEM with 25ng/ml supGM-CSF is added. All cells that adhere to the surface of the plate are considered alveolar macrophages (AM) as previously determined by Chen *et al*. (PMID:3288696). After new media is added, AMs are stimulated with 200ng/ml LPS (Sigma, L2630-10MG). Harvest supernatant 6hrs post-stimulation. Harvested supernatant was sent to Eve technologies for cytokine analysis. Statistics were performed using GraphPad prism.

### Cytotoxicity Assay

Cytotoxicity was assessed by measuring the release of LDH into the media (LDH-Cytotoxicity Colorimetric Assay Kit II; BioVision) according to the manufacturer’s protocol.

## Results

### Production of a murine GM-CSF-secreting HEK-293T cell line

A murine GM-CSF (mGM-CSF) containing plasmid (Addgene, 74465) was used as a template to amplify mGM-CSF, which we inserted into a plasmid with a pSico backbone (SFig1, Supplemental File). The vector contains a bidirectional EF1a promoter driving puromycin resistance gene and mCherry on one side and mGM-CSF on the other (SFig1B, Fig1A). Lentivirus containing mGM-CSF, or a control construct was generated and transduced into HEK293Ts and monitored using flow cytometry (SFig2). Post puromycin selection for 7 days, our control and mGM-CSF constructs were incorporated into the genomes of HEK293T cells (at a rate of >85%) (Fig1B). In order to determine how much GM-CSF was being produced by the HEK293T line we plated and grew 2×10^6^ cells for 3 days, isolated the supernatant and performed an ELISA to measure the concentration which was found to be 181.7ng/ml (Extended Methods). The cell line was subsequently frozen down into aliquots and thawed to assess stability. Protein secretion was assessed again, by ELISA, and the stock secreted mGM-CSF at 181.9±ng/ml (Fig1C). There was no significant difference between the concentration of mGM-CSF in the supernatant of the initial or the freeze thawed HEK293T cell line indicating that the line is stable.

**Figure 1:**
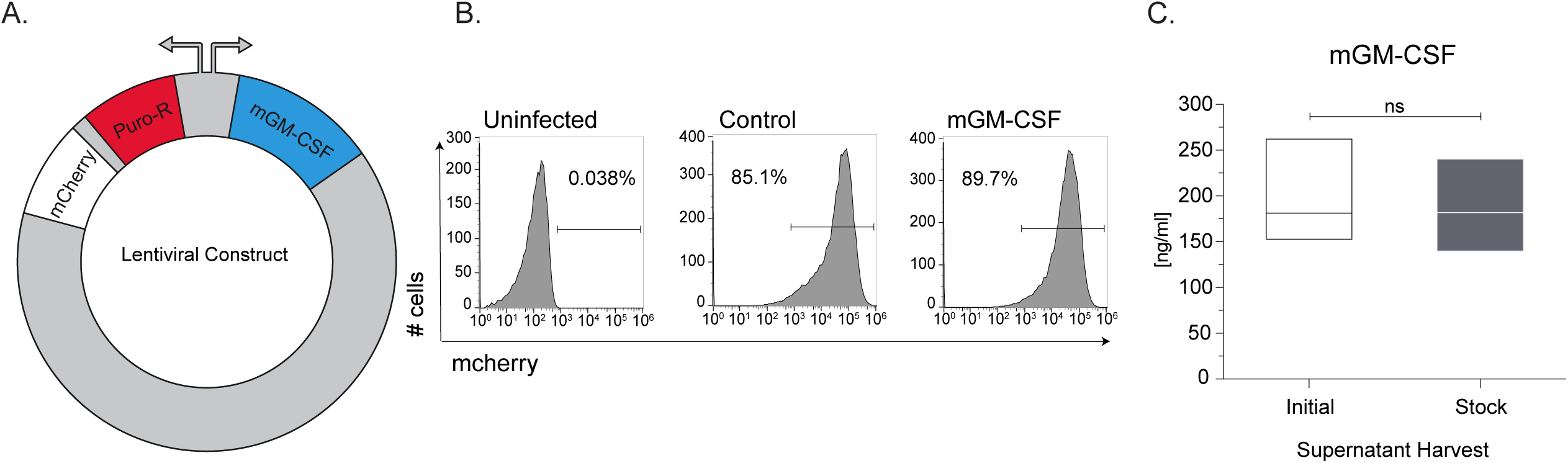
Generation of mouse GM-CSF secreting HEK293T cell line. **(A)** Schematic of lentiviral plasmid construct used to generate the stable HEK293T mGM-CSF expressing cell line. **(B)** Uninfected, Control plasmid and mGM-CSF plasmid lentivirally infected HEK239T cells were puromycin selected for mCherry expression and assessed using flow cytometry. **(C)** Protein secretion and cell-line stability of mGM-CSF HEK293T cells was confirmed ELISA, using initial and stock cell-line. Error bars represent the standard deviation of biological triplicates. Student’s t-tests were performed using GraphPad Prism. (SN) indicates not significantly different.

### Bone-Marrow Cells differentiated with supGM-CSF yield a higher number of Dendritic Cells compared to pGM-CSF

To assess the efficacy of the mGM-CSF-rich supernatant (supGM-CSF), bone marrow (BM) cells were treated with 10ng/ml or 25 ng/ml of supGM-CSF and compared to the same concentrations of a commercially obtained purified GM-CSF (pGM-CSF) (*11–13*). As a control for differentiation, we also generated bone-marrow derived macrophages (BMDMs) using supM-CSF obtained from L929 cells (*14*) (Fig2A).

**Figure 2:**
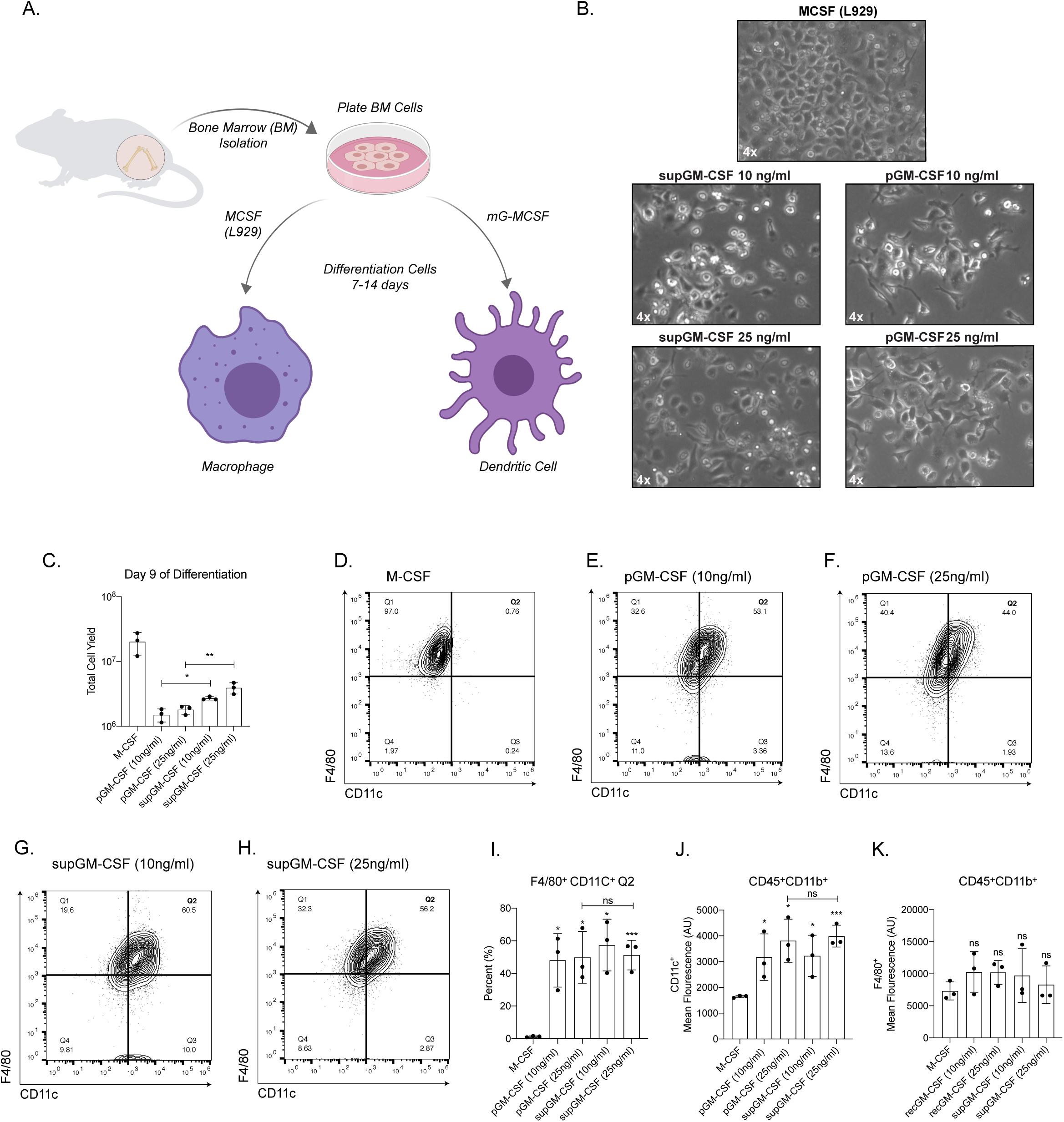
Generation of primary Dendritic Cells using purified GM-CSF or supernatant GM-CSF. **(A)** Schematic of differentiation of primary bone marrow-derived macrophages (BMMs) or primary bone marrow-derived dendritic cells (BMDCs). **(B)** Images of primary BMMs and BMDCs under a light microscope after 7 days of differentiation. **(C)** Total number of live cells generated from using one of 5 conditions: M-CSF, purified GM-CSF (pGM-CSF) at 10ng/ml, pGM-CSF at 25ng/ml, supernatant GM-CSF (supGM-CSF) at 10ng/ml or supGM-CSF at 25ng/ml. **(D)** Gating of CD45+ and CD11b+ M-CSF differentiated cells are represented in a quadrant contour plot to determine the expression of F4/80 and CD11c. Same gating strategy is used for **(E)** pGM-CSF at 10ng/ml, **(F)** pGM-CSF at 25ng/ml, **(G)** supGM-CSF at 10ng/ml and **(H)** supGM-CSF at 25ng/ml. **(I)** Graphical representation for the percentage of M-CSF or GM-CSF differentiated cells that are F4/80+ and CD11c+ (Q2). (J) Graphical representation for the mean fluorescence of CD11c+. (K) Graphical representation for the mean fluorescence of F4/80+. Error bars represent the standard deviation of n=5. Student’s t-tests were performed using GraphPad Prism. Asterisks indicate statistically significant differences between mouse lines (*p ≥ 0.05, **p ≥ 0.01, ***p ≥ 0.005).

Cells incubated with M-CSF, pGM-CSF and supGM-CSF were fully differentiated by day 7 and could be cultured until day 14 (Fig2A). Morphologically, BM differentiated with supGM-CSF and pGM-CSF, at either 10ng/ml or 25ng/ml, have a more stellate morphology as is expected in dendritic cells compared to M-CSF differentiated macrophage cells. There is no morphological difference between pGM-CSF or supGM-CSF differentiated cells, at either 10ng/ml or 25ng/ml (Fig 2B). At day 9 of differentiation, cells were harvested and counted. When comparing the pGM-CSF to the supGM-CSF, supGM-CSF yields significantly more viable cells in comparison to pGM-CSF (Fig 2C). After culture with M-CSF, pGM-CSF or supGM-CSF, cells were assessed for purity by flow cytometry based on previously published panels (*15, 16*). After gating on Live+/CD45+/CD11b+ cells purity was determined based on the proportion of the population expressing CD11c and F4/80, shown as a quadrant (SFig3, Fig2D-H). Percent of cells in quadrant 2 (F4/80+ and CD11c+) was significantly higher for all GM-CSF treated cells, in comparison to M-CSF treated cells, as is expected since CD11c is a dendritic cell marker. However, pGM-CSF and supGM-CSF were not significantly different (Fig2 I). As expected, the mean fluorescence intensity for dendritic cell marker CD11c+ cells was significantly higher for all GM-CSF treated cells in comparison to M-CSF (Fig2 J), while F4/80+ expression was equal for every treatment (Fig2 K). While supGM-CSF is able to generate more viable cells than pGM-CSF, there is no significant difference in cellular purity between supGM-CSF and pGM-CSF (both at ∼85% purity).

### BMDCs generated using supGM-CSF function as efficiently as BMDCs generated using pGM-CSF

Dendritic cells are the professional antigen presenting cells of the immune system. In order to test the ability of our supGM-CSF to generate functional DCs we measured their levels of MHC-Class II expression using flow cytometry (*17*). M-CSF differentiated cells express little MHC class II, while pGM-CSF and supGM-CSF express robust levels of MHC class II, when differentiated with 10ng/ml or 25ng/ml concentration of GM-CSF (Fig3A-B, SupFig3A-B). Dendritic cells differentiated with 10ng/ml GM-CSF show statistically higher mean fluorescence intensity for MHC class II for supGM-CSF in comparison to pGM-CSF (SFig.4 C). Interestingly, dendritic cells differentiated with 25ng/ml of either pGM-CSF or supGM-CSF exhibit no significant difference in MHC class II protein expression when evaluated by mean fluorescence intensity (Fig.3 C). In addition to assessing MHC Class II levels, we also assessed the ability of the pGM-CSF and supGM-CSF BMDCs to respond to inflammatory stimulation. Both pGM-CSF and supGM-CSF BMDCs were stimulated with LPS (200ng/ml) for the indicated time course and IL6 mRNA expression was measured by quantitative PCR (qPCR) (Fig.3 C). Thus, determining that the supGM-CSF derived cells maintain their inflammatory activity. Finally, AIM2 inflammasome activity was measured through IL6 and IL1b protein secretion of both purified and supernatant GM-CSF. Here we observe that supGM-CSF produces significantly higher expression of IL6 and IL1b (Fig.3 D-E) suggesting that these dendritic cells are more sensitive to inflammatory activation compared to DCs generated using pGM-CSF.

**Figure 3:**
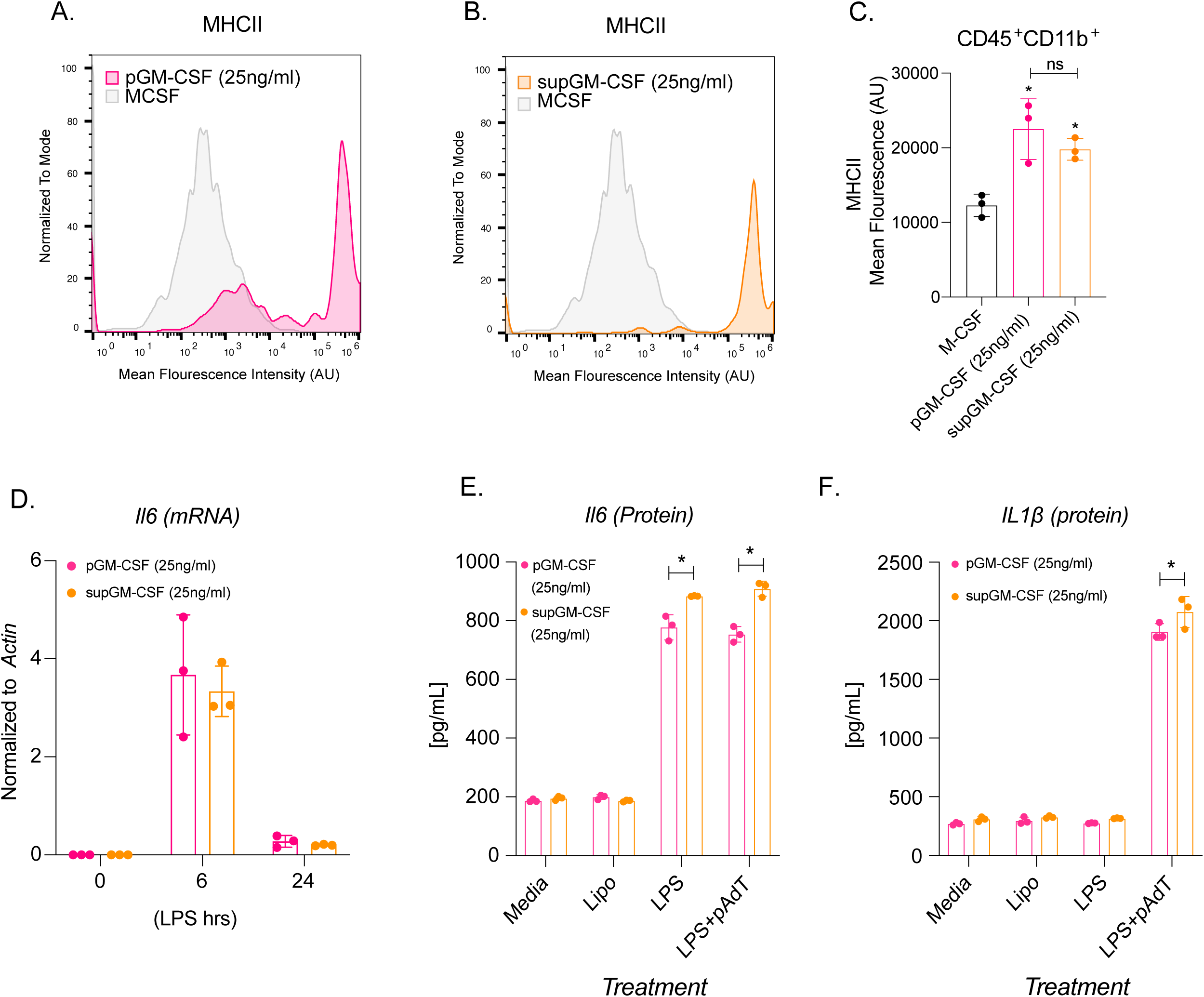
Dendritic Cells differentiated with supGM-CSF retain Dendritic Cell activity and are more inflammatory in comparison to pGM-CSF. **(A)** Histogram overlay of CD45+ and CD11b+ M-CSF (grey) and pGM-CSF (pink) or **(B)** supGM-CSF (orange) differentiated cells expressing MHC II. **(C)** Graphical representation of MHC II mean fluorescence intensity (MFI) of M-CSF, pGM-CSF, or supGM-CSF differentiated cells. **(D)** Il6 transcript measure by RTq-PCR from pGM-CSF (pink) or supGM-CSF (orange) DCs stimulated with LPS for 0, 6, and 24hrs. (E) Dendritic cells differentiated with pGM-CSF (pink) or sup-GM-CSF (orange) were treated with lipofectamine, LPS (24hrs) or LPS (2hrs) and polydA:dT (6hrs) secreted IL-6 or IL-1β **(E)** was measured by ELISA. Student’s t-tests were performed using GraphPad Prism. Asterisks indicate statistically significant differences between mouse lines (*p ≥ 0.05, **p ≥ 0.01, ***p ≥ 0.005).

### supGM-CSF can maintain viable and inflammatory inducible Alveolar Macrophages

mGM-CSF is a critical protein factor that is not only necessary for driving primary dendritic cell differentiation, but also for maintaining primary alveolar macrophages (AM) in culture (*18, 19*). To test the ability of supGM-CSF to maintain AM in culture, we harvested bronchiolar lavage fluid (BALF) from 10 WT wild-type mice. We pooled these cells, counted and then plated them in 25ng/ml supGM-CSF supplemented media. 24 h post-harvest, using a light microscope, we tested that AMs attached (Fig. 4). Viability of AMs were assessed by measuring the amount of lactate dehydrogenase (LDH) in the media. The amount of measured LDH correlates directly with the cell number lysed (*20*). Our data indicate that supGM-CSF does not negatively affect the cell culture, while the controls indicate there are healthy cultured cells not resistant to apoptosis (Fig.4 B). AMs were stimulated with LPS for 24 hr and cytokines were measured by ELISA. Inflammatory inducible proteins are significantly upregulated in AMs cultured in supGM-CSF media, including proteins IL6, TNFa, MDC, MCP-1, IP10, KC and Rantes (Fig.4 C-I). supGM-CSF cultured primary alveolar macrophages maintain their pro-inflammatory activation programming.

**Figure 4:**
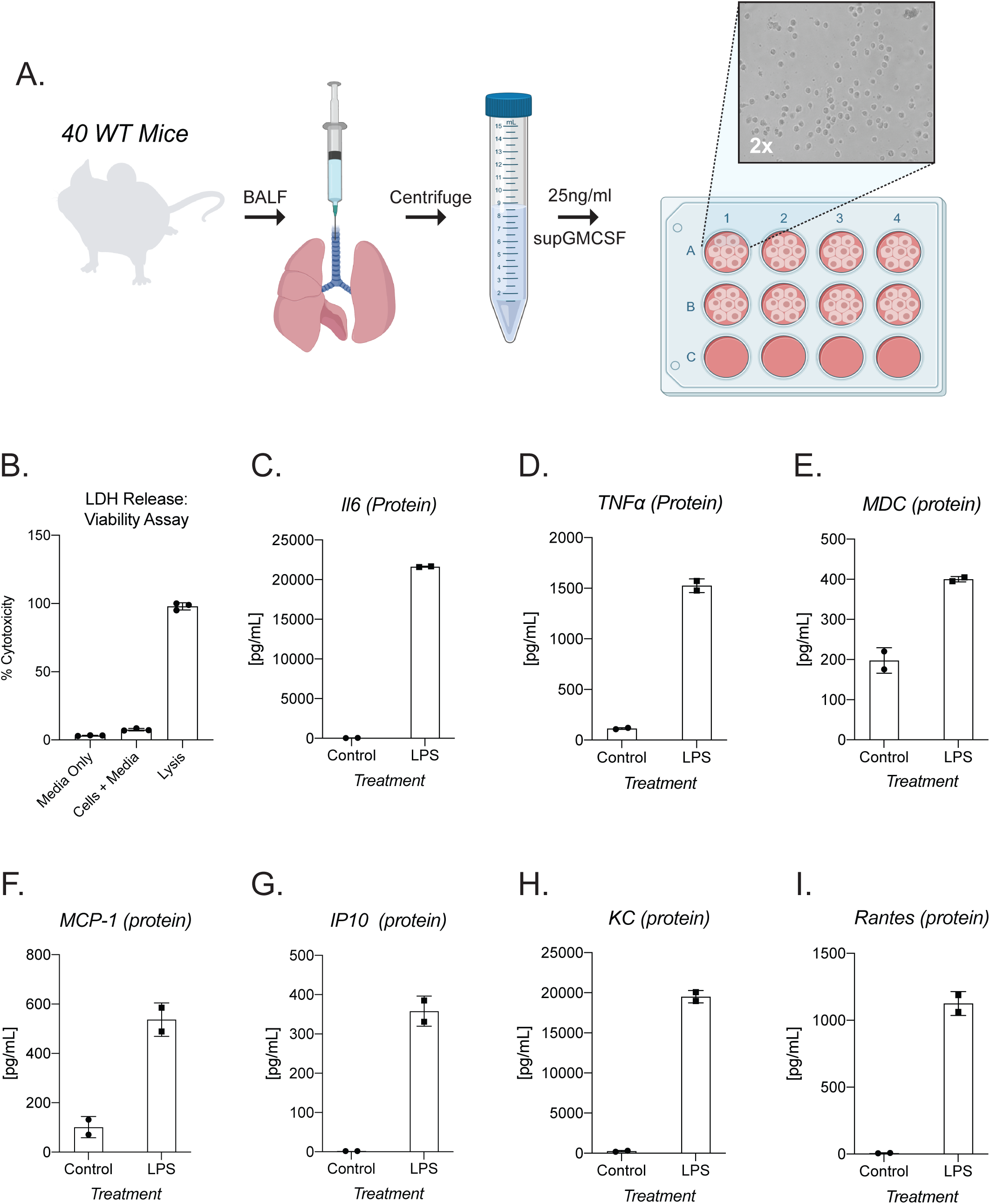
supGM-CSF can differentiate functional Alveolar Macrophages. **(A)** Schematic of alveolar macrophage supGM-CSF harvesting and differentiating experiment. **(B)** The cellular viability of differentiated alveolar macrophages was measured by LDH, where media, cells, and lysis of cells were measured. **(C)** Secreted IL6 protein was measured from supGM-CSF differentiated alveolar macrophages treated with and without LPS for 6hrs, as well as **(D)** TNFa, **(E)** MDC, **(F)** MCP-1, **(G)** IP10, **(H)** KC and **(I)** Rantes. Student’s t-tests were performed using GraphPad Prism. Asterisks indicate statistically significant differences between mouse lines (*p ≥ 0.05, **p ≥ 0.01, ***p ≥ 0.005).

## Discussion

Dendritic cells and macrophages have been cultured and studied for the last 40 years, leading to many advances in culturing protocols. Flt3L is often utilized as a factor for BMDC differentiation, it is now appreciated that Flt3L DCs are representative of steady-state resident DCs, while GM-CSF BMDCs mirror the transcriptional programing of pro-inflammatory recruited cells (*21, 22*). Granulocyte-macrophage colony-stimulating factor (GM-CSF), differentiated bone marrow cells are widely used as a model system for conventional DC development (*23, 24*), as well as sustaining primary alveolar macrophages in culture (*25*). Current strategies for the generation of murine BMDCs or culturing of primary alveolar macrophages utilize store bought purified recombinant GM-CSF or a stable cell-line expressing recGM-CSF called J558L (*10*). J558L, a murine GM-CSF, secreting cell-line is utilized for both BMDC and alveolar macrophage culturing, however it is an immune cell that also secretes IL-10 which can alter the transcriptional programing of the cells when culturing and only secretes 80ng/ml (*10, 18, 19*).

Using supernatant from GM-CSF secreting HEK293T cells, can serve as an alternative to purified GM-CSF for murine BMDCs or maintenance of primary alveolar macrophages. We have successfully cloned and stably integrated murine GM-CSF into HEK293T cells, which consistently secrete 180ng/ml of GM-CSF (Fig. 1). Bone marrow differentiated with our supGM-CSF produces more cells by day 9 compared to pGM-CSF, but both GM-CSF sources generate an equal percentage of pure dendritic cells based on previously published gating strategies (*15, 16*) (Fig1C and I). Additionally, using more GM-CSF also will produce more viable and proliferating cells, which can be further enhanced with higher concentrations of supGM-CSF (Fig2C). Commercial GM-CSF can be used instead of our HEK293T supGM-CSF, but it is expensive, and GM-CSF has to be added every 2 days to cells during the differentiation process. Our supGM-CSF is much more cost effective compared to purified GM-CSF from a number of companies (Supplemental Table 1). From one harvest of supGM-CSF you can generate ∼10 ug, which will be the cost of 50ml of complete media (Extended Methods), while 10 ug of purified GM-CSF from Thermo Fisher will cost $276. The purity of BMDCs generated from pGM-CSF and supGM-CSF does not differ when assessed by flow cytometry based on previously published panels (*15, 16*) (SFig3, Fig2D-H).

DCs are professional antigen presenting cells and therefore express high levels of MHC Class II (*21, 26*). BMDCs differentiated with either Flt3L or GM-CSF have comparable T cell activation, and therefore MHC II expression (*22*). BMDCs differentiated with either pGM-CSF or supGM-CSF express MHC-II at a comparable level, when measured by mean fluorescence through flow cytometry (Fig3A-C).

As previously stated, picking a cell line to express a recombinant protein is very important. If an immune cell is chosen, one risks the chance of having immune factors being secreted in the supernatant, leading to priming and activation of either pro- or anti-inflammatory transcriptional programs (*10, 18, 19*). By performing a time-course stimulation or overnight LPS stimulation, our study indicates supGM-CSF does not inhibit the inflammatory program transcriptionally when measuring the transcript or protein level of IL6 (Fig 3D-E). supGM-CSF and pGM-CSF differentiated BMDCs both retain the ability to activate the inflammasome pathway, leading to the secretion of IL1b (Fig3D-F), while this pathway is now contested in DCs our results for GM-CSF is true (*27*).

Not only does our study provide data to support the use of supGM-CSF compared to pGM-CSF for BMDCs, but we also show that supGM-CSF can be utilized to sustain the culturing of primary alveolar macrophages from murine bronchoalveolar lavage fluid (BALF) (*10, 18*). After 48 h of supGM-CSF cultured BALF, the adhered alveolar macrophages (AMs) were >95% viable (Fig4B). More importantly, when stimulating the cells with LPS overnight cytokines such as IL6, TNFa, KC and Rantes were all inducible (Fig. 4C-I). Thus, supGM-CSF maintains AMs in culture and does not inhibit the pro-inflammatory programming of the immune cells.

Taken together our results show that a pure BMDC population and inflammatory inducible DCs or AMs can be established by culturing BM cells or BALF with a crude supernatant from our GM-CSF HEK293T cell line. This line will provide the research community with a more cost-effective alternative to commercially available GM-CSF.

## Author Contributions

SCo and SCa has designed the study and cell line. EKR has analyzed data for all figures. SZ has performed all flow cytometry for figures 1, 2 and 3. SZ has characterized the cell line and generated extended methods. EKR has personally performed experiments for data used in Figure 3 and 4 and supplemental figure 3 and table. EKR has written and edited manuscript. SCa, SCo, SZ have edited manuscript. SCa has provided necessary funding for project.

## Data and Materials Availability

Our vector maps are available in supplemental materials.

## Figure Legends

**Supplemental Figure 1:**
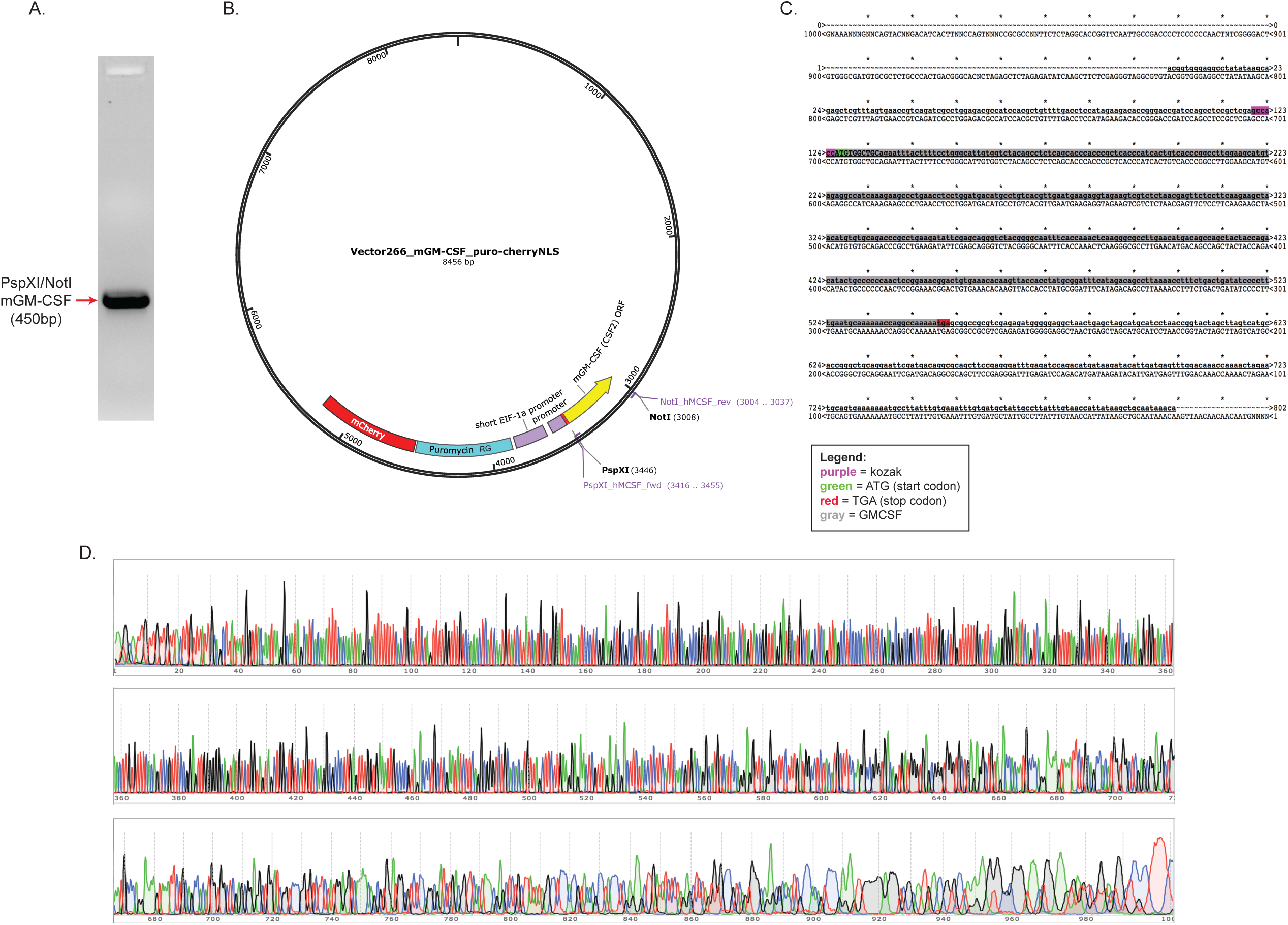
Validation of cloned murine GM-CSF. **(A)** Image of PspXI and NotI digested PCR product of murine GM-CSF. **(B)** Detailed 266 vector map of mGM-CSF, restriction sites, promoter, antibiotic selection and mCherry sequence. **(C)** Alignment of Sanger sequencing results of mGM-CSF and 266 Vector map strategy. **(D)** Quality of sanger sequencing results of murine GM-CSF.

**Supplemental Figure 2:**
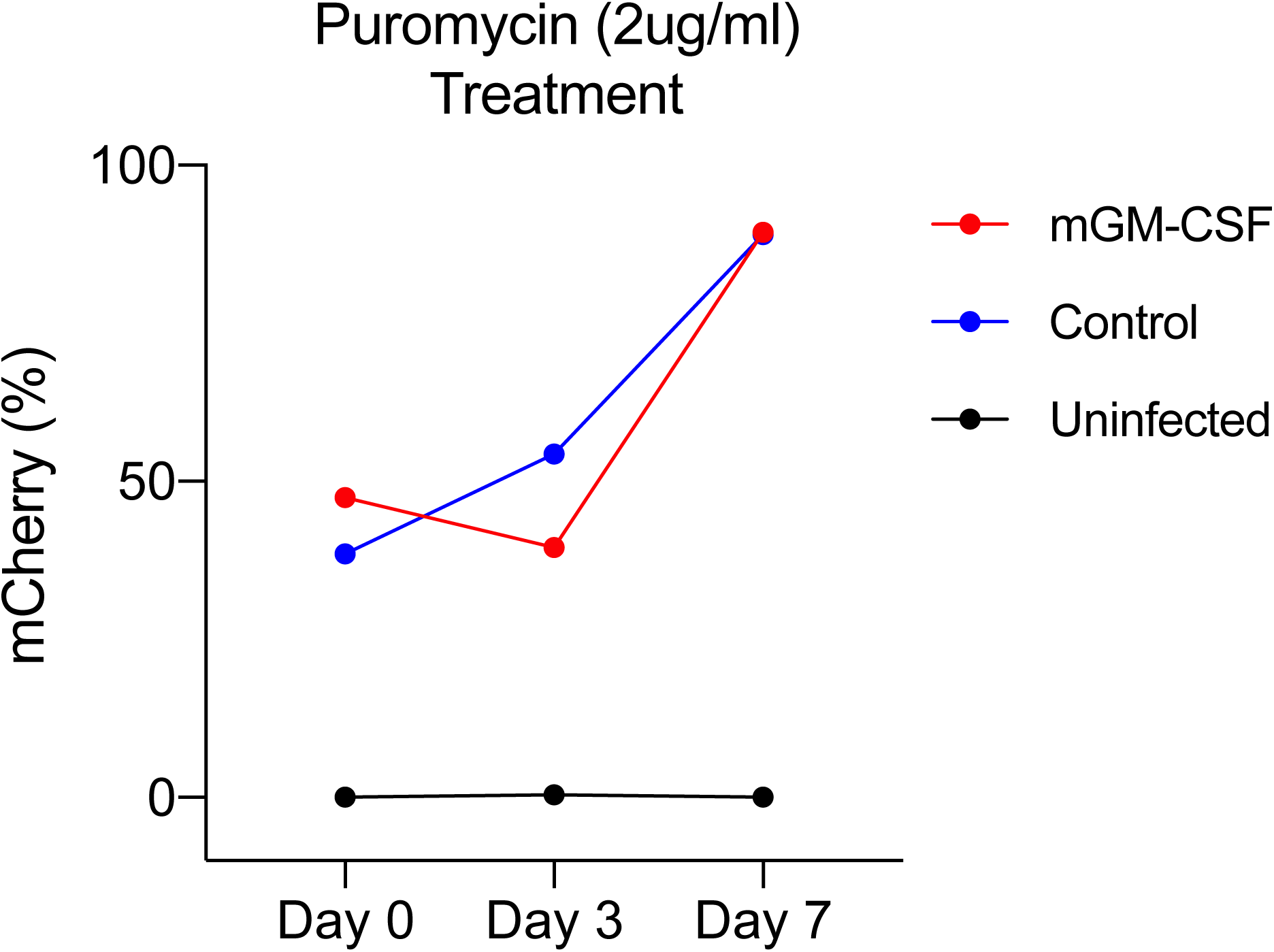
mCherry expression in HEK293T cell line. HEK-293T cell line lentiviral integration of GM-CSF was selected using puromycin (2ug/ml) over a week. Flow cytometry was used to assess fluorescence intensity.

**Supplemental Figure 3:**
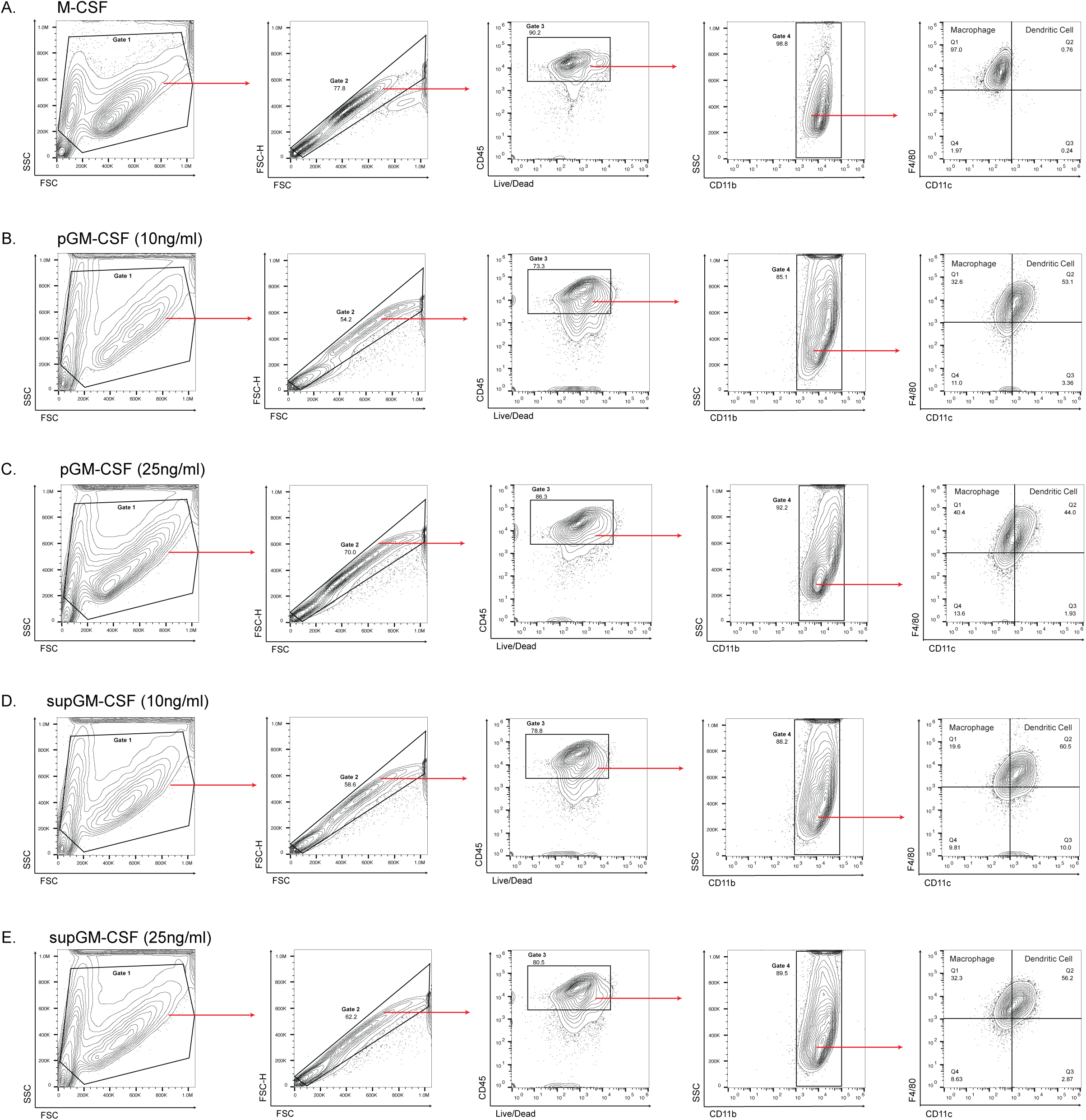
Gating strategy for identifying dendritic cell and macrophage populations. M-CSF and GM-CSF differentiated cells were both put through the same gating strategy. The 5 gating plots are for the **(A)** M-CSF, **(B)** pGM-CSF (10ng/ml), **(C)** pGM-CSF (25ng/ml), **(D)** supGM-CSF (10ng/ml), **(E)** supGM-CSF (25ng/ml) differentiated cells. Non-debris cells were gated for in gate 1, then singlets in gate 2, followed by CD45+ and live cells were gated for in gate 3, then CD11b+ cells were gated in gate 4 finally this population of cells were visualized using F4/80+ and CD11c+ markers and gate were put into quadrants.

**Supplemental Figure 4:**
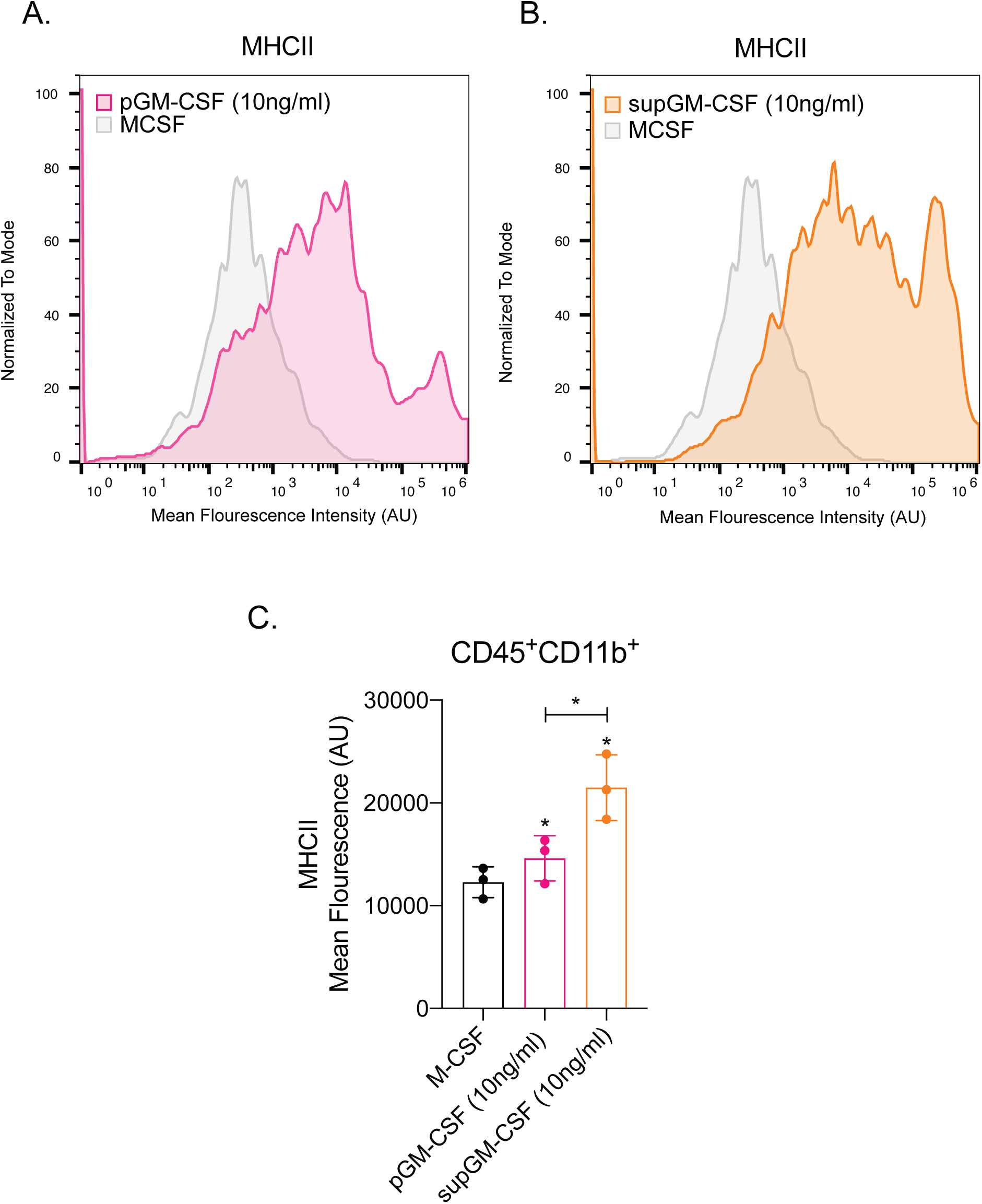
Expression of MHC II on dendritic cell populations differentiated with 10ng/ml pGM-CSF or supGM-CSF. **(A)** Histogram overlay of CD45+ and CD11b+ M-CSF (grey) and pGM-CSF (pink) or **(B)** supGM-CSF (orange) differentiated cells expressing MHC II. **(C)** Graphical representation of MHC II mean fluorescence of M-CSF, pGM-CSF, or supGM-CSF differentiated cells. Student’s t-tests were performed using GraphPad Prism. Asterisks indicate statistically significant differences between mouse lines (*p ≥ 0.05, **p ≥ 0.01, ***p ≥ 0.005).

## Extended Methods

Comprehensive Methods in the Creation of an mGM-CSF-Producing HEK293T Cell Line, Collection of mGM-CSF-rich Supernatant, and Using Cell Line Supernatant to Generate Dendritic Cells From Myeloid Progenitors

### Creating Cell Line

1. Acquire mGM-CSF gene (Addgene: Plasmid #74465)
  a. Sequence to check for correct composition
2. PCR mGM-CSF gene
  a. Create and acquire primers with required restriction sites (NotI-HF and PspXI)
  b. PCR
3. Purify/isolate mGM-CSF gene (band size?)
4. Ligation of mGM-CSF gene with 681 bidirectional vector plasmid
  a. Restriction digest 681 bidirectional vector with Not1-HF and PspXI and purify
  b. Restriction digest modified mGM-CSF gene with Not1-HF and PspXI and purify
  c. Ligate vector with gene
5. Transform E. coli. with 681 + GM-CSF plasmid construct
6. Extract and isolate 681 + GM-CSF plasmid construct
  a. Incubate E. coli in liquid LB
  b. Miniprep
  c. Colony PCR and run through the gel to determine sample with GM-CSF gene inserted in 681 vector
7. Create lentiviral constructs containing plasmid
8. Infect HEK293 cells using lentiviral constructs
9. Select for cells with 681 + GM-CSF plasmid construct
  a. Allow cells to recuperate from infection and grow to confluence
  b. Puro-select
  c. FACS to determine concentration of mCherry+ cells (>90% required)

### Collecting mGM-CSF-rich supernatant

1. Allow for HEK293 cells with plasmid construct to grow to confluency
2. (To be redone in T-175) Plate 2 million cells per 10 cm plate
3. Incubate for 3 days
4. Collect supernatant
5. If needed to confirm the concentration of mGM-CSF in the supernatant, perform ELISA (∼200 ng/ml)

### *In a T-175 flask

1. Allow for HEK293 cells with plasmid construct to grow to confluency
2. Plate 9 million cells in 50 ml DMEM per T-175 flask
3. Incubate for 3 days
4. Collect supernatant
5. If needed to confirm the concentration of mGM-CSF in the supernatant, perform ELISA (∼200 ng/ml)

### Generation of Bone-Marrow-Derived DCs

*Day 0*

1. Sacrifice one or two mice and reserve femur and tibia of mice
2. Extract bone marrow cells from femur and tibia
3. In 6 well plate, plate equal amounts of cells into each well
  a. Total media should be equal to 2 ml per well (cells + DMEM + cytokine)
4. Incubate for three days *Day 3*
5. On third day of differentiation, wash off dead cells
  a. Aspirate media
  b. Perform a rough PBS wash
  c. Aspirate PBS
  d. Replace media
  *Day 4 (if cells reach confluency)*
6. On fourth day of differentiation, move cells onto 10 cm plate
  a. Save conditioned media
  b. Perform normal PBS wash?
  c. Aspirate PBS
  d. Using cell scraper, gently scrape cells off plate
  e. Transfer cells onto 10 cm plate
  f. Replace media
    i. Total media should be 10 ml (cells + DMEM + cytokine)
7. Incubate for 2-3 days *Day 5 or 6*
8. Replace media
  a. Save conditioned media
  b. Perform normal PBS wash?
  c. Aspirate PBS
  d. Replace media (10 ml total; 6.750 ml DMEM + 2 ml conditioned media + ∼1.250 ml supernatant)
9. Incubate for 1-2 days *Day 7 or until usage before day 14*
10. Replace media every 2-3 days
11. Cells finished differentiation. Ready for use.

---

**Supplemental Table 1:**
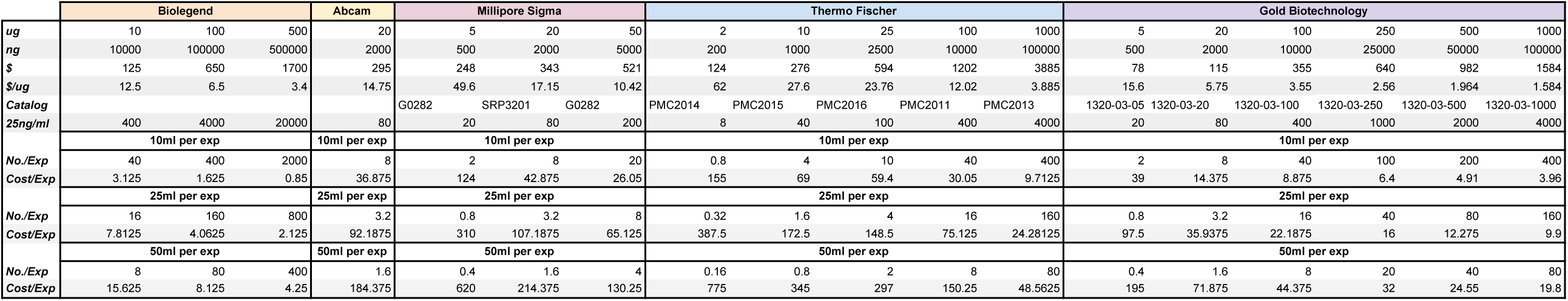
Cost of purified murine GM-CSF.

|  | Biologend |  |  | Abcam | Millipore Sigma |  |  | Thermo Fischer |  |  |  |  | Gold Biotechnology |  |  |  |  |  |
| --- | --- | --- | --- | --- | --- | --- | --- | --- | --- | --- | --- | --- | --- | --- | --- | --- | --- | --- |
| *ug* | 10 | 100 | 500 | 20 | 5 | 20 | 50 | 2 | 10 | 25 | 100 | 1000 | 5 | 20 | 100 | 250 | 500 | 1000 |
| *ng* | 10000 | 100000 | 500000 | 2000 | 500 | 2000 | 5000 | 200 | 1000 | 2500 | 10000 | 100000 | 500 | 2000 | 10000 | 25000 | 50000 | 100000 |
| *\$* | 125 | 650 | 1700 | 295 | 248 | 343 | 521 | 124 | 276 | 594 | 1202 | 3885 | 78 | 115 | 355 | 640 | 982 | 1584 |
| *\$/ug* | 12.5 | 6.5 | 3.4 | 14.75 | 49.6 | 17.15 | 10.42 | 62 | 27.6 | 23.76 | 12.02 | 3.885 | 15.6 | 5.75 | 3.55 | 2.56 | 1.964 | 1.584 |
| *Catalog* |  |  |  |  | G0282 | SRP3201 | G0282 | PMC2014 | PMC2015 | PMC2016 | PMC2011 | PMC2013 | 1320-03-05 | 1320-03-20 | 1320-03-100 | 1320-03-250 | 1320-03-500 | 1320-03-1000 |
| *25ng/ml* | 400 | 4000 | 20000 | 80 | 20 | 80 | 200 | 8 | 40 | 100 | 400 | 4000 | 20 | 80 | 400 | 1000 | 2000 | 4000 |
|  | 10ml per exp |  |  | 10ml per exp | 10ml per exp |  |  | 10ml per exp |  |  |  |  | 10ml per exp |  |  |  |  |  |
| *No./Exp* | 40 | 400 | 2000 | 8 | 2 | 8 | 20 | 0.8 | 4 | 10 | 40 | 400 | 2 | 8 | 40 | 100 | 200 | 400 |
| *Cost/Exp* | 3.125 | 1.625 | 0.85 | 36.875 | 124 | 42.875 | 26.05 | 155 | 69 | 59.4 | 30.05 | 9.7125 | 39 | 14.375 | 8.875 | 6.4 | 4.91 | 3.96 |
|  | 25ml per exp |  |  | 25ml per exp | 25ml per exp |  |  | 25ml per exp |  |  |  |  | 25ml per exp |  |  |  |  |  |
| *No./Exp* | 16 | 160 | 800 | 3.2 | 0.8 | 3.2 | 8 | 0.32 | 1.6 | 4 | 16 | 160 | 0.8 | 3.2 | 16 | 40 | 80 | 160 |
| *Cost/Exp* | 7.8125 | 4.0625 | 2.125 | 92.1875 | 310 | 107.1875 | 65.125 | 387.5 | 172.5 | 148.5 | 75.125 | 24.28125 | 97.5 | 35.9375 | 22.1875 | 16 | 12.275 | 9.9 |
|  | 50ml per exp |  |  | 50ml per exp | 50ml per exp |  |  | 50ml per exp |  |  |  |  | 50ml per exp |  |  |  |  |  |
| *No./Exp* | 8 | 80 | 400 | 1.6 | 0.4 | 1.6 | 4 | 0.16 | 0.8 | 2 | 8 | 80 | 0.4 | 1.6 | 8 | 20 | 40 | 80 |
| *Cost/Exp* | 15.625 | 8.125 | 4.25 | 184.375 | 620 | 214.375 | 130.25 | 775 | 345 | 297 | 150.25 | 48.5625 | 195 | 71.875 | 44.375 | 32 | 24.55 | 19.8 |

